# The *Gossypium stocksii* genome as a novel resource for cotton improvement

**DOI:** 10.1101/2021.02.23.432605

**Authors:** Corrinne E. Grover, Daojun Yuan, Mark A. Arick, Emma R. Miller, Guanjing Hu, Daniel G. Peterson, Jonathan F. Wendel, Joshua A. Udall

## Abstract

Cotton is an important textile crop whose gains in production over the last century have been challenged by various diseases. Because many modern cultivars are susceptible to several pests and pathogens, breeding efforts have included attempts to introgress wild, naturally resistant germplasm into elite lines. *Gossypium stocksii* is a wild cotton species native to Africa, which is part of a clade of vastly understudied species. Most of what is known about this species comes from pest resistance surveys and/or breeding efforts, which suggests that *G. stocksii* could be a valuable reservoir of natural pest resistance. Here we present a high-quality *de novo* genome sequence for *G. stocksii*. We compare the *G. stocksii* genome with resequencing data from a closely related, understudied species (*G. somalense*) to generate insight into the relatedness of these cotton species. Finally, we discuss the utility of the *G. stocksii* genome for understanding pest resistance in cotton, particularly resistance to cotton leaf curl virus.

## Introduction

The cotton genus, *Gossypium,* is responsible for providing the majority of natural textile fiber through the cultivation of its four domesticated species. While most research and resource development is devoted to the two major polyploid crop species, i.e., *G. hirsutum* and *G. barbadense*, the cultivated diploid species *G. herbaceum* and *G. arboreum* comprise a significant share of the cotton market in certain countries (Wendel *et al.* 1989; Basu 1996; Guo *et al.* 2006; Khadi *et al.* 2010; Kranthi 2018). Native to Africa, these latter two species are nestled within a clade of additional African species that possess short non-spinnable fiber, but which may be valuable as sources of various disease and/or stress resistant traits (Yik and Birchfield 1984; Rudgers *et al.* 2004; Nazeer *et al.* 2014; Rahman *et al.* 2017).

*Gossypium stocksii,* is a diploid cotton species native to Eastern Africa whose subsection, *Pseudopambak* (E-genome cottons (Wang *et al.* 2018)), is thought to be earliest diverging lineage in the African clade (Wendel & Grover, 2015). E-genome cottons, including *G. stocksii* (E_1_), *G. somalense* (E_2_), *G. areysianum* (E_3_), and *G. incanum* (E_4_), may be sources of valuable traits including disease resistance. While both *G. stocksii* and *G. somalense* have resistance to reniform nematode (Yik and Birchfield 1984), only *G. stocksii* has reported resistance to cotton leaf curl disease (CLCuD) (Nazeer *et al.* 2014). Spread by white flies (Briddon and Markham 2000), the virus that causes CLCuD can have devastating effects on crop yield, as exhibited by the Pakistan epidemic in the early 1990s (Rahman *et al.* 2017), which resulted in massive financial losses over the course of five years. By some estimates, CLCuD is capable of decreasing total yield up to 90%, yet none of the major *G. hirsutum* cultivars exhibit resistance (Mammadov *et al.* 2018).

Because *G. stocksii* germplasm may be a useful source of resistance traits, interspecific material derived from crosses between *G. stocksii* and the commercially important *G. hirsutum* have been evaluated for a number of traits, including resistance to CLCuD and possible improvements in fiber. Research has shown that the F1 generation of a doubled *G. stocksii* x *G. hirsutum* cross not only has resistance to CLCuD, but also exhibits increased fiber strength relative to the parents (Nazeer *et al.* 2014). More recently, comparisons among hexaploid hybrids derived from crosses between *G. hirsutum* and other wild diploid species suggests that four wild diploid species, including *G. stocksii*, are potentially valuable for fiber breeding programs (Konan *et al.* 2020).

Although there has been interest in *G. stocksii* for breeding purposes, genomic resources are virtually non-existent for this species. Here we describe a high-quality, *de novo* genome sequence for *G. stocksii,* a valuable source of disease resistance in cotton and a potential source for improving fiber in domesticated cotton.

## Methods & Materials

### Plant material and sequencing methods

Mature leaves from *G. stocksii* (E_1_ grown under greenhouse conditions at Brigham Young University (BYU) were collected for PacBio sequencing. A CTAB-based method was used to extract high-quality DNA (Kidwell and Osborn 1992), which was quantified on a Qubit Fluorometer (ThermoFisher, Inc.). A BluePippen instrument (Sage Science, LLC) was used to size-select for only fragments >18 kb, as verified using a Fragment Analyzer (Advanced Analytical Technologies, Inc). Size-selected DNA was sent to the BYU DNA Sequencing Center (DNASC) for PacBio (Pacific Biosciences) library construction and sequencing on a total of 20 PacBio cells. Canu V1.6 was used to assemble the raw sequencing reads using default parameters (Koren *et al.* 2017).

Young leaf tissue was also used for DNA extraction and HiC library construction (Belton *et al.* 2012) by PhaseGenomics LLC. These HiC libraries were sequenced on an Illumina HiSeq 2500 (2×125 bp) at the BYU DNASC. Resulting HiC reads were used to join contigs, and the association frequency between paired-ends was used to correct the assembly using JuiceBox (Durand et al., 2016). The final genome sequence of *G. stocksii* was generated via a custom python script available through PhaseGenomics LLC, yielding 13 assembled chromosomes.

### Repeat and gene annotation

Transposable elements were annotated using a combination of RepeatMasker (Smit *et al.* 2015) and “One code to find them all” (Bailly-Bechet *et al.* 2014). A custom library based off of Repbase 23.04 (Bao *et al.* 2015) combined with cotton-specific repeats (Grover *et al.* 2020) was used in conjunction with RepeatMasker to mark repeats in the genome. Adjacent matches were merged using “One code to find them all,” and the output was aggregated and summarized in R/4.0.3 (R Core Team 2017) using *dplyr* /0.8.1 (Wickham *et al.* 2015). All code is available at https://github.com/Wendellab/stocksii.

The *G. stocksii* genome was annotated using existing RNA-seq data from various tissues of closely related species (Supplementary Table 1). Specifically, the following tissues were used: *Gossypium arboreum* developing seeds and seedling (SRR617075, SRR617073, SRR617068, SRR617067, SRR959508), *Gossypium davidsonii* roots and leaves (SRR2132267), *Gossypium herbaceum* seed and developing fiber (SRR959585, SRR10675236, SRR10675235, SRR10675234, SRR10675237), *Gossypium longicalyx* leaf, stem, and flower (SRR1174182, SRR1174179, SRR6327757, SRR6327758, SRR6327759), *Gossypium raimondii* leaf, seed, stem, petal, meristem, and floral tissues (SRR617009, SRR617011, SRR617013, SRR8267554, SRR8267566, SRR8878565, SRR8878526, SRR8878661, SRR8878800, SRR8878534, SRR8878745), *Gossypium thurberi* leaf, root, and stem (SRR8267623, SRR8267616, SRR8267619), and *Gossypium trilobum* leaf, root, and stem (SRR8267606, SRR8267582, SRR8267601). Each library was downloaded from the Short Read Archive (SRA), and all RNA-seq data was mapped to the hard-masked *G. stocksii* genome using hisat2 [v2.1.0] (Kim *et al.* 2015). BRAKER2 [v2.1.2] (Hoff *et al.* 2019) was trained with GeneMark [v4.38] (Borodovsky and Lomsadze 2011) generated annotations which were also used to train Augustus [v3.3.2] (Stanke *et al.* 2006). StringTie [v2.1.1] (Pertea *et al.* 2015) and Cufflinks [v2.2.1] (Ghosh and Chan 2016) generated *de novo* RNA-seq assemblies were combined with a Trinity [v2.8.6] (Grabherr *et al.* 2011) reference-guided assembly and splice junction information from Portcullis [v1.2.2] (Mapleson *et al.* 2018) in Mikado [v1.2.4] (Venturini *et al.* 2018). MAKER2 [v2.31.10] (Holt and Yandell 2011; Campbell *et al.* 2014) was used to integrate gene predictions from (1) BRAKER2 trained Augustus, (2) GeneMark, and (3) Mikado, also using evidence from all *Gossypium* ESTs available from NCBI (nucleotide database filtered on “txid3633” and “is_est”) and a database composed of all curated proteins in Uniprot Swissprot [v2019_07] (UniProt Consortium 2008) combined with the annotated proteins from the *G. hirsutum* (https://www.cottongen.org/species/Gossypium_hirsutum/jgi-AD1_genome_v1.1) and *G. raimondii* (Paterson *et al.* 2012) genomes. SNAP [v2013-02-16] and Augustus were trained with the predicted annotations from Maker. Maker was run a second time with the newly trained Augustus and SNAP models, along with the other inputs from the first iterations. Annotation edit distance (AED - (Eilbeck *et al.* 2009; Holt and Yandell 2011; Yandell and Ence 2012) was used to score each gene model relative to EST and protein evidence, and gene models with an AED less than 0.35 were retained. Gene models were functionally annotated using InterProScan [v5.47-82.0] (Jones *et al.* 2014) and BlastP [v2.9.0+] (Camacho *et al.* 2009) searches against the Uniprot SwissProt database. Orthologous relationships between *G. stocksii* and other diploid cottons were determined via OrthoFinder (Emms and Kelly 2015, 2019). Proteins from *G. longicalyx (Grover et al. 2020)*, *G. arboreum* (Li *et al.* 2014; Du *et al.* 2018; Huang *et al.* 2020), *G. herbaceum (Huang et al. 2020), G. raimondii* (Paterson *et al.* 2012; Udall *et al.* 2019a), *G. turneri (Udall et al. 2019a)*, and *G. australe (Cai et al. 2019)* were downloaded from CottonGen (https://www.cottongen.org; (Yu *et al.* 2014) and run using default parameters. Code is available from https://github.com/Wendellab/stocksii.

### Comparison to *G. somalense*

Three DNA libraries of *G. somalense* (E_2_), a close relative of *G. stocksii* (SRA; SRR3560160-SRR3560162), were used to provide a preliminary comparison of the two species. Raw reads were mapped to the newly generated *G. stocksii* genome using the Spack (Gamblin *et al.* 2015) implementation of bwa v0.7.17-rgxh5dw (Li and Durbin 2009). Single-nucleotide polymorphisms (SNPs) in *G. somalense* were called relative to *G. stocksii* using the Sentieon pipeline (Kendig *et al.* 2019) (Spack version sentieon-genomics/201808.01-opfuvzr), which is an optimization of existing methods, such as GATK (McKenna *et al.* 2010). This pipeline included read deduplication, indel realignment, and genotyping. The three libraries represent technical replicates of the *G. somalense* sequencing, and were therefore merged after read deduplication. Parameters for mapping and SNP calling follow standard practices, and are available in detail at https://github.com/Wendellab/stocksii. The resulting variant file was filtered for read depth using vcftools (Spack version version 0.1.14-v5mvhea) (Danecek *et al.* 2011), only retaining sites with a minimum of 10 reads and a maximum of 100 reads. GenomeTools (Gremme *et al.* 2013) was used to convert the annotation file to gtf format, which was used in conjunction with SnpEff (Cingolani *et al.* 2012) to annotate and predict the effects of the SNP differences between *G. stocksii* and *G. somalense*.

Divergence between the two species was estimated using--window-pi from vcftools in 100 kb, non-overlapping windows, which estimated the average number of differences per window. Diversity was parsed by region by first intersecting the filtered VCF with the relevant feature (e.g., exon, intron, etc.) from the *G. stocksii* annotation using intersectBed from bedtools2 (Spack version 2.27.1-s2mtpsu) (Quinlan 2014) to get a list of SNP sites associated with that region. The original, filtered VCF was then used in conjunction with vcftools--window-pi and the flag--positions, which limits the analysis to only the specified sites (e.g., exon, intron, intergenic). Diversity/divergence results were parsed in R/4.0.3 using dplyr (Wickham *et al.* 2015) and plotted using ggplot2 (Wickham 2016). Relevant code and detailed pipeline analysis can be found at https://github.com/Wendellab/stocksii.

To provide a comparative framework for qualitative interpretation of the amount of divergence between *G. stocksii* and *G. somalense,* two other species pairs (i.e., *G. herbaceum-G. arboreum* and *G. raimondii-G. gossypioides,* were also subjected to SNP calling/filtering and calculation of π in 100 kb windows, as outlined above for *G. stocksii-G. somalense.* Here, the genome of *G. herbaceum* (Huang *et al.* 2020) was used as a reference for *G. arboreum* reads (SRR8979980; (Page *et al.* 2013), and *G. raimondii* (Udall *et al.* 2019a) was used as a reference for reads from *G. gossypioides* (SRR3560148 and SRR3560149). Genomes and annotations were both downloaded from CottonGen (Yu *et al.* 2014).

### Data availability

The *G. stocksii* genome sequence is available at NCBI under PRJNA701967 and through CottonGen (https://www.cottongen.org/). Raw data is available from the SRA under PRJNA701967. Supplemental files are available from figshare.

## Results and Discussion

### Genome assembly and annotation

We report a high-quality *de novo* genome sequence for *G. stocksii* covering 93% of the 1531 Mb genome (Hendrix and Stewart 2005). PacBio reads (58X coverage) were initially assembled into 316 contigs with an N50 of 17.8 Mb. These contigs were then ordered and oriented using both HiC and Bionano evidence to produce a chromosome level assembly (n=13) with an average length of 110 Mb (1424 Mb total) and containing only 5.7 kb of gap sequence across all chromosomes. BUSCO (Waterhouse *et al.* 2017) analysis of the genome (Table 1) indicates a general completeness with only 4.2% of BUSCOs either fragmented (0.9%) or missing (1.5%). Over 97% complete BUSCOs were recovered, most of which were single copy (88.9%, versus 8.7% duplicated). The LTR Assembly Index (LAI) (Ou *et al.* 2018) was also within guidelines for “reference-quality” genomes (LAI = 10-20; *G. stocksii* LAI = 15.4), and dotplots (Figure 1) with existing high-quality cotton genome assemblies (Paterson *et al.* 2012; Du *et al.* 2018; Udall *et al.* 2019a, 2019b; Grover *et al.* 2020; Huang *et al.* 2020) further indicates the high quality nature of this genome.

**Table 1:** BUSCO results for the genome and annotation.

| Table 1: BUSCO results for the genome and annotation. |  |  |
| --- | --- | --- |
|  | Genome | Annotation |
| Complete BUSCOs (C) | 2271 (97.6%) | 2227 (95.8%) |
| Complete and single-copy BUSCOs (S) | 2068 (88.9%) | 1888 (81.2%) |
| Complete and duplicated BUSCOs (D) | 203 (8.7%) | 339 (14.6%) |
| Fragmented BUSCOs (F) | 20 (0.9%) | 26 (1.1%) |
| Missing BUSCOs (M) | 35 (1.5%) | 73 (3.1%) |
| Total BUSCO groups searched | 2326 |  |

**Figure 1.**
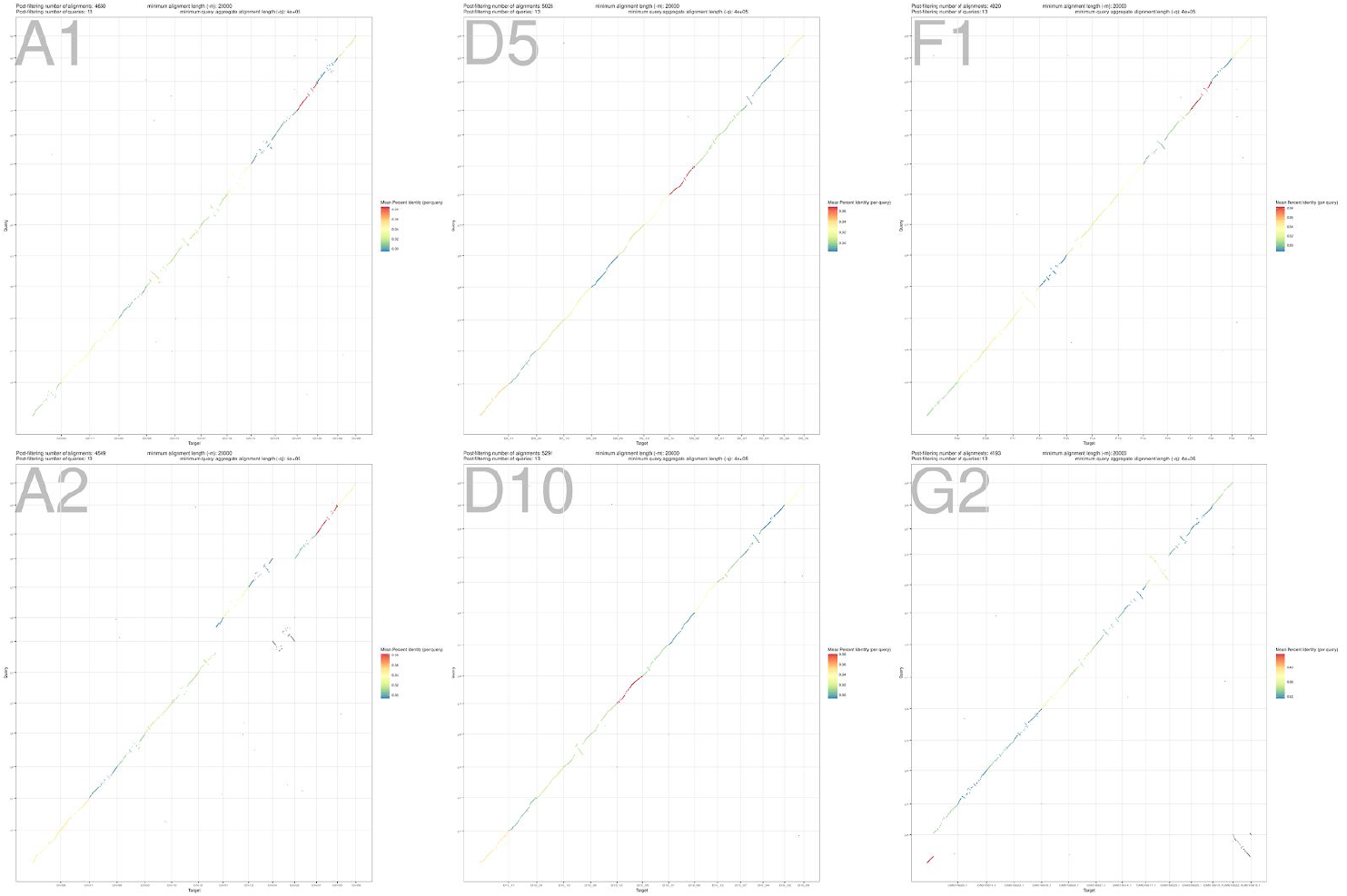
Pairwise comparisons of *G. stocksii* with *G. herbaceum* (A1; (Huang *et al.* 2020), *G. raimondii* (D5; (Udall *et al.* 2019a), *G. longicalyx* (F1; (Grover *et al.* 2020), *G. arboreum* (A2; (Huang *et al.* 2020), *G. turneri* (D10; (Udall *et al.* 2019a), and *G. australe* (G2; (Cai *et al.* 2019).

Annotation of the *G. stocksii* genome revealed 37,889 transcripts representing 34,928 unique genes, similar to other cotton diploid genomes (range 37,505 to 43,952; (Paterson *et al.* 2012; Du *et al.* 2018; Udall *et al.* 2019a; Grover *et al.* 2020; Huang *et al.* 2020). BUSCO analysis of the annotation (Table 1) exhibits recovery, similar to the whole genome BUSCO. Ortholog analysis between *G. stocksii* and these previously published cotton diploids produces 23,399 orthogroups (Supplementary File 1) containing at least one *G. stocksii* gene (range 18,785 in *G. australe* to 27,913 in *G. arboreum;* (Huang *et al.* 2020), comprising 68.5% of the total orthogroups. Notably, five species-specific orthogroups were recovered containing a total of 67 genes (Table 2), 62 of which are argonaute-like proteins (Supplementary Table 2). On average, over half of the transcripts (22,403) are placed in a simple 1:1 relationship in pairwise comparisons between *G. stocksii* and another cotton diploid genome (Supplementary Table 3).

**Table 2:**
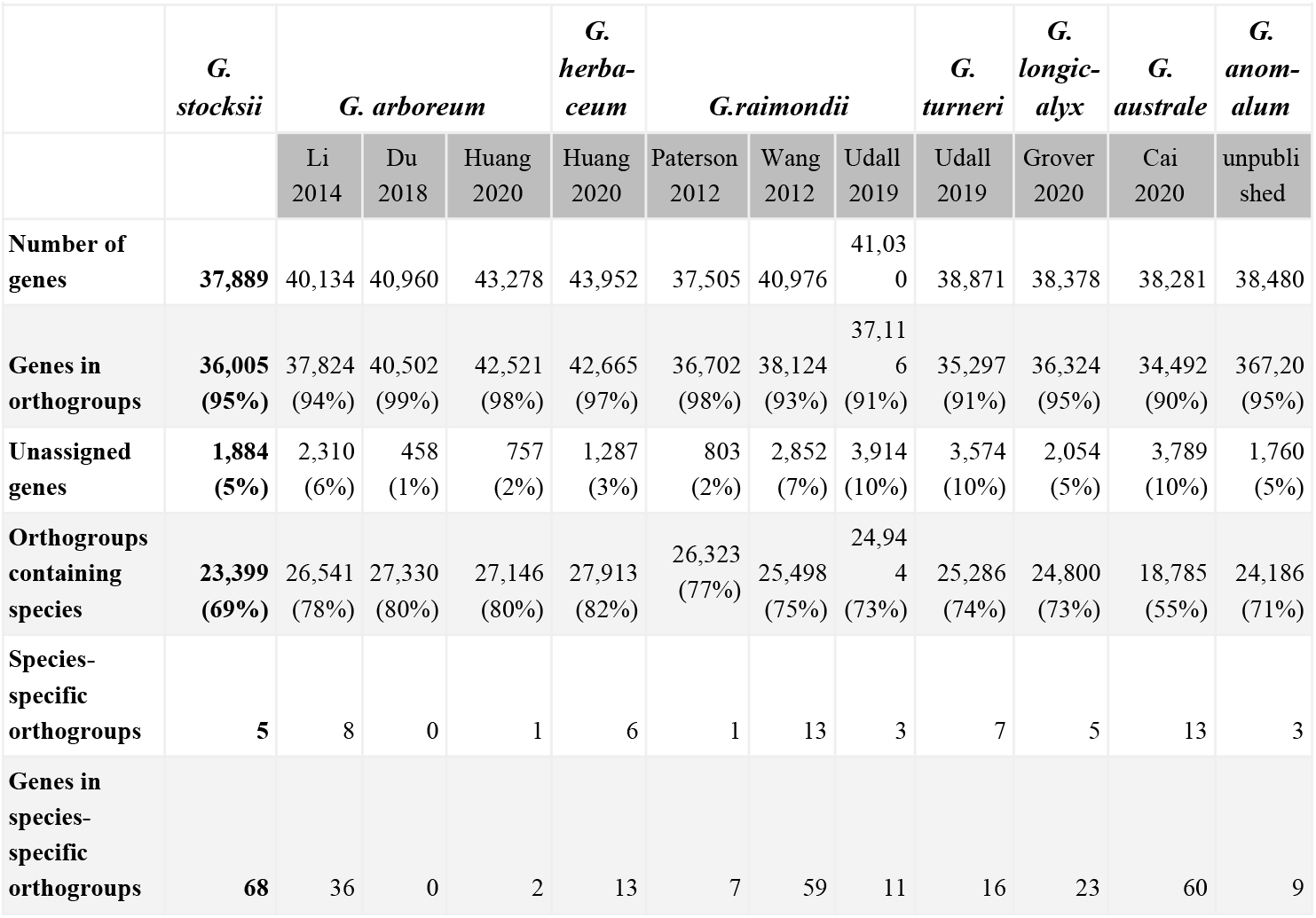
Orthogroup relationships between *G. stocksii* and other cotton diploid genomes.

|  | *G.*<br>*stocksii* | *G. arboreum* |  |  | *G.*<br>*herba-*<br>*ceum* | *G. raimondii* |  |  | *G.*<br>*turneri* | *G.*<br>*longic-*<br>*alyx* | *G.*<br>*australe* | *G.*<br>*anom-*<br>*alum* |
| --- | --- | --- | --- | --- | --- | --- | --- | --- | --- | --- | --- | --- |
|  |  | Li<br>2014 | Du<br>2018 | Huang<br>2020 | Huang<br>2020 | Paterson<br>2012 | Wang<br>2012 | Udall<br>2019 | Udall<br>2019 | Grover<br>2020 | Cai<br>2020 | unpubli<br>shed |
| **Number of**<br>**genes** | **37,889** | 40,134 | 40,960 | 43,278 | 43,952 | 37,505 | 40,976 | 41,030 | 38,871 | 38,378 | 38,281 | 38,480 |
| **Genes in**<br>**orthogroups** | **36,005**<br>**(95%)** | 37,824<br>(94%) | 40,502<br>(99%) | 42,521<br>(98%) | 42,665<br>(97%) | 36,702<br>(98%) | 38,124<br>(93%) | 37,116<br>(91%) | 35,297<br>(91%) | 36,324<br>(95%) | 34,492<br>(90%) | 367,20<br>(95%) |
| **Unassigned**<br>**genes** | **1,884**<br>**(5%)** | 2,310<br>(6%) | 458<br>(1%) | 757<br>(2%) | 1,287<br>(3%) | 803<br>(2%) | 2,852<br>(7%) | 3,914<br>(10%) | 3,574<br>(10%) | 2,054<br>(5%) | 3,789<br>(10%) | 1,760<br>(5%) |
| **Orthogroups**<br>**containing**<br>**species** | **23,399**<br>**(69%)** | 26,541<br>(78%) | 27,330<br>(80%) | 27,146<br>(80%) | 27,913<br>(82%) | 26,323<br>(77%) | 25,498<br>(75%) | 24,944<br>(73%) | 25,286<br>(74%) | 24,800<br>(73%) | 18,785<br>(55%) | 24,186<br>(71%) |
| **Species-**<br>**specific**<br>**orthogroups** | **5** | 8 | 0 | 1 | 6 | 1 | 13 | 3 | 7 | 5 | 13 | 3 |
| **Genes in**<br>**species-**<br>**specific**<br>**orthogroups** | **68** | 36 | 0 | 2 | 13 | 7 | 59 | 11 | 16 | 23 | 60 | 9 |

Transposable element (TE) content was assessed by *de novo* TE prediction via RepeatMasker (Bailly-Bechet *et al.* 2014; Smit *et al.* 2015), indicating that repeats occupy approximately 43% of the 1531 Mbp genome (Table 3). Consistent with other plant genomes, *Ty3/gypsy* predominate the *G. stocksii* genome, comprising over 90% of the detected repetitive elements. Ty1/*’copia* elements and DNA elements (as a whole) were substantially less represented, accounting for only 43 and 13 Mb, respectively, in the present analysis.

**Table 3:** Repeat types and predicted copy numbers in the *G. stocksii* genome

| Element type | Fragments | Copies | SoloLTR | Total_Mb |
| --- | --- | --- | --- | --- |
| **DNA** | 15,047 | 9,190 | 0 | 13.37 |
| DNA/EnSpmCACTA | 1,138 | 682 | 0 | 2.02 |
| DNA/Harbinger | 1 | 1 | 0 | 0.00 |
| DNA/hAT | 1,858 | 1195 | 0 | 0.84 |
| DNA/hAT-Tip100 | 18 | 11 | 0 | 0.02 |
| DNA/L1 | 923 | 461 | 0 | 1.08 |
| DNA/MarinerTc1 | 71 | 39 | 0 | 0.05 |
| DNA/MuDR | 11,030 | 6,797 | 0 | 9.36 |
| DNA/MULE-MuDR | 6 | 3 | 0 | 0.00 |
| DNA/PIF-Harbinger | 2 | 1 | 0 | 0.00 |
| **LTR** | 906,998 | 487,232 | 269,000 | 650.36 |
| LTR | 34 | 33 | 0 | 0.00 |
| LTR/Copia | 45,806 | 27,117 | 9,567 | 42.96 |
| LTR/Gypsy | 861,158 | 460,082 | 259,433 | 607.39 |
| **Total** | 922,045 | 496,422 | 0 | 663.73 |

### Comparison of *G. stocksii* with *G. somalense*

*Gossypium stocksii* is part of a clade of approximately seven species (subsection Pseudopambak), but relatively little is known about the members of this subsection, including questions regarding species circumscription and the possibility of unrecognized taxa (Fryxell 1979, 1992; Vollesen 1987). A comparison between *Gossypium stocksii* and the closely related *G. somalense* reveals considerable divergence between these two species, with 39.7 M interspecific SNPs evenly distributed among the 13 chromosomes (Table 4). As expected, most of the variation (94%, or 37.1 M SNPs) is found in the intergenic space, only 30% of which is found near genes (±5 kb up- or down-stream). An assessment of nucleotide distance between *G. stocksii* and *G. somalense* (here measured as π in VCFtools) reveals a modest distance between these two species (mean π=0.0116; 100 kb windows) that is intermediate between the very closely related sister species *G. arboreum* and *G. herbaceum* (Renny-Byfield *et al.* 2016; Huang *et al.* 2020) and the more distantly related species *G. gossypioides* and *G. raimondii* (subgenus *Houzingenia*; (Grover *et al.* 2019). On a per-chromosome basis, the pairwise *G. stocksii-G.somalense* π estimates range from an average of 0.0098 on chromosome E05 to 0.0126 on chromosome E03 (Figure 2).

**Table 4:** Comparison of *G. somalense* resequencing with the *G. stocksii* genome

| Chromosome | Length | Number of variants | Variant rate | Average pi |
| --- | --- | --- | --- | --- |
| E01 | 116,888,287 | 3,307,654 | 2.83% | 0.0124 |
| E02 | 88,432,363 | 2,545,417 | 2.88% | 0.0120 |
| E03 | 125,861,959 | 3,604,952 | 2.86% | 0.0126 |
| E04 | 106,357,252 | 3,070,741 | 2.89% | 0.0124 |
| E05 | 106,784,523 | 2,598,022 | 2.43% | 0.0098 |
| E06 | 118,523,473 | 3,312,902 | 2.80% | 0.0120 |
| E07 | 94,613,334 | 2,855,058 | 3.02% | 0.0118 |
| E08 | 122,663,403 | 3,390,477 | 2.76% | 0.0117 |
| E09 | 82,831,125 | 2,285,194 | 2.76% | 0.0106 |
| E10 | 114,014,049 | 3,343,978 | 2.93% | 0.0120 |
| E11 | 117,232,604 | 3,111,632 | 2.65% | 0.0104 |
| E12 | 114,623,165 | 3,068,541 | 2.68% | 0.0112 |
| E13 | 115,591,314 | 3,227,954 | 2.79% | 0.0121 |
| **Total** | **1,424,416,851** | **39,722,522** | 2.79% | 0.0116 |
| **Total (genic)** | **128,641,547** | **2,578,738** | **2.00%** | 0.0007 |
|  | SNP location | Number of variants | Proportion of variants |  |
|  | Intergenic | 37,188,781 | 93.62% |  |
|  | upstream | 5,461,093 | 13.75% |  |
|  | downstream | 5,121,580 | 12.89% |  |
|  |  |  | 0.00% |  |
|  | Exon | 831,986 | 2.09% |  |
|  | missense | 471,909 | 1.19% |  |
|  | silent | 352,203 | 0.89% |  |
|  | nonsense | 14,780 | 0.04% |  |
|  | Intron | 1,746,752 | 4.40% |  |
|  | UTR, 5' | 63,862 | 0.16% |  |
|  | UTR, 3' | 71,297 | 0.18% |  |

**Figure 2.**
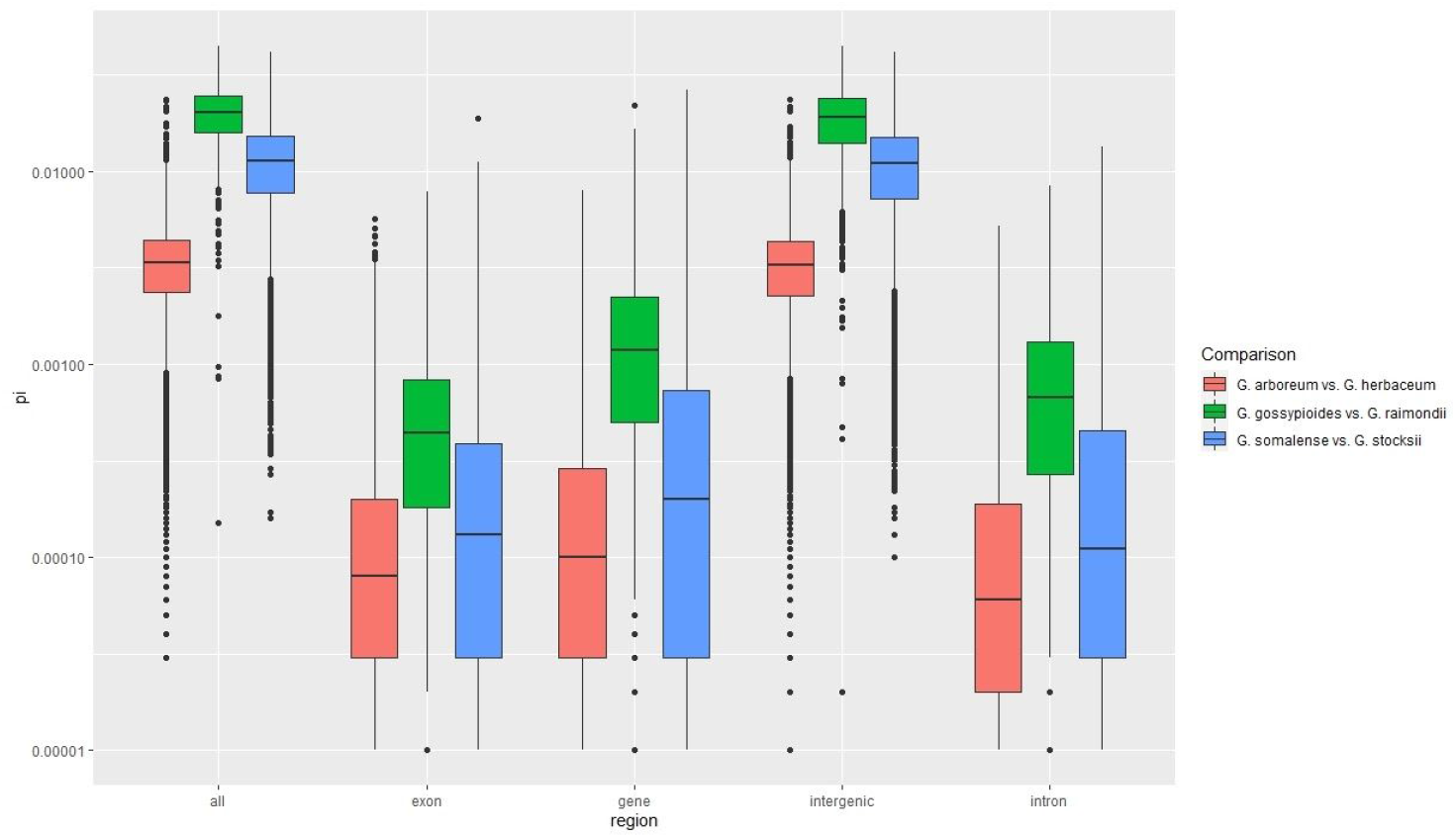
Pairwise comparisons of π for *G. somalense* and *G. stocksii,* with *G. arboreum* versus *G. herbaceum* and *G. gossypioides* versus *G. raimondii* for comparison. Here, π is calculated individually for each species pair for the entire dataset (all) and for the specified subset of SNPs (i.e., exonic, genic, intragenic, and intronic). Colors reflect the individual comparisons, with lines to represent the mean and points to represent outliers. Because π is calculated between two samples each, the values here reflect the pairwise divergence between samples.

Although genic regions have far fewer SNPs, SNPs in these regions still account for 2.6 M of the 39.7 M total (Table 3). Intron-based SNPs outweigh exon-based SNPs in a 2:1 ratio, accounting for 4.4% and 2.1% of the overall SNPs, respectively. Most exon-based SNPs are minimally disruptive, either conferring silent (352,203) or missense (471,909) changes (Table 3); very few (14,780) produced predicted nonsense changes. Similar to other species pairs in *Gossypium,* the average nucleotide distance in genes was far lower than the overall distance (0.007 versus 0.0116, respectively), indicating a close relationship between these two species in their gene space. Given that *G. somalense* does not exhibit the sample level of resistance to CLCuD (Nazeer *et al.* 2014; Anjum *et al.* 2015), but does show other forms of pest resistance (Yik and Birchfield 1984; Shim *et al.* 2018), future comparisons including multiple accessions of both species may shed insight into the evolution of natural pest resistance in cotton species.

### *G. stocksii* as a resource for disease resistance

Whereas domesticated varieties of *G. hirsutum* are highly susceptible to CLCuD (Rehman *et al.* 2017), *G. stocksii* exhibits natural resistance (Nazeer *et al.* 2014). The molecular basis of CLCuD resistance in cotton is not well understood (Rahman *et al.* 2017), although genetic analyses indicate that CLCuD resistance is likely controlled by one or few dominant genes with possible epistatic modifiers (Knight 1948; Ali 1997; Haidar *et al.* 2003; Rahman *et al.* 2005; Ahuja *et al.* 2007), thereby making it a prime target for breeding programs and/or genetic modification. While the success of CRISPR/Cas9 in controlling similar viral diseases and the continued lack of success in controlling CLCuD using conventional methods (Iqbal *et al.* 2016) has piqued interest in genome modification enhancing resistance, little research has focused on the genomic basis of CLCuD resistance.

Preliminary research in a CLCuD-resistant accession of *G. arboreum* identified 1,062 differentially expressed genes (DEG) between challenged and unchallenged plants (Naqvi *et al.* 2017), 17 of which were considered prime candidates for conferring disease resistance. Of those 17 genes, 16 were placed in orthogroups that also contained one or more *G. stocksii* homologs (Table 5), with the sole exception of the gene putatively encoding ‘phytosulfokines 3’ (i.e., Cotton_A_25246_BGI-A2_v1.0), which plays a role in pathogen response in lotus (Wang *et al.* 2015). Most orthogroups were comparable in size between the *G. arboreum* genome used to detect DEG and our *G. stocksii* annotation, aside from OG0000284 (the cysteine protease ervatamin-B like genes), which was composed of five tandemly arrayed genes in *G. arboreum,* but only two in *G. stocksii*; the relevance of these genes to CLCuD defense is unclear. The largest orthogroup that contained one of the top DEG candidates was orthogroup OG0000074, which is composed of resistance gene (i.e., R-gene) analogs (Naqvi *et al.* 2017); notably, *G. stocksii* appears to have one additional copy of this gene. Similarity at the protein level between the *G. arboreum* DEG and its closest *G. stocksii* homolog is generally high (i.e., 95%, on average), although it drops as low as 73.4% in the poorly conserved ervatamin-B like orthogroup (Table 5). These results indicate that similar genes may operate in CLCuD resistance in *G. stocksii*; however, comparative expression data from infected and uninfected plants is required to understand whether the two species use similar pathways to avoid infection by the CLC virus.

**Table 5:** Ortholog identification for previously identified CLCuD candidates and copy numbers in other published genomes. Numbers in parentheses indicate the number(s) of genes from that orthogroup that are genomically clustered. Orthogroups without parenthetical notation indicate all members from that species are genomically dispersed.

| CLCuD candidate* | Total genes | Average per genome | Closest match | protein similarity | Orthogroup | *G. arboreum*<br>Li *et al.* 2014 | *G. stocksii* | *G. herbaceum*<br>Huang *et al.* 2020 | *G. arboreum*<br>Du *et al.* 2018 | *G. arboreum*<br>Huang *et al.* 2020 | *G. raimondii*<br>Wang *et al.* 2012 | *G. raimondii*<br>Paterson *et al.* 2012 | *G. raimondii*<br>Udall *et al.* 2019 | *G. turneri*<br>Udall *et al.* 2019 | *G. longicalyx*<br>Grover *et al.* 2020 | *G. anomalum*<br>unpublished | *G. australe*<br>Cai *et al.* 2020 |
| --- | --- | --- | --- | --- | --- | --- | --- | --- | --- | --- | --- | --- | --- | --- | --- | --- | --- |
| Cotton_A_03097_BGI-A2_v1.0 | 140 | 11.7 | Gosto.013G220600-1 | 88.6% | OG0000074 | 8 (6+1+1) | 9 (2 + 1 + 1 + 4) | 12 (2 + 1 + 1 + 8) | 15 (1 + 1 + 12) | 12 (1 + 1 + 10) | 12 (2 + 1 + 1 + 8) | 6 (6) | 9 (1 + 8) | 31 (31) | 15 (1 + 1 + 13) | 10 (1 + 9) | 1 |
| Cotton_A_22786_BGI-A2_v1.0 | 63 | 5.3 | Gosto.005G342100-1 | 73.4% | OG0000284 | 5 (5) | 2 (2) | 5 (5) | 5 (5) | 5 (5) | 6 (6) | 7 (7) | 6 (6) | 7 (7) | 6 (6) | 6 (6) | 3 (3) |
| Cotton_A_34570_BGI-A2_v1.0 | 43 | 3.6 | Gosto.006G064800-1 | 87.6% | OG0000679 | 3 | 4 (1 + 3) | 5 (1 + 4) | 5 (1 + 3 + 1) | 5 (1 + 4) | 4 (1 + 1 + 2) | 5 (3 + 2) | 2 | 4 (1 + 3) | 2 | 3 | 1 |
| Cotton_A_07057_BGI-A2_v1.0 | 43 | 3.6 | Gosto.005G102100-1 | 100.0% | OG0000687 | 4 | 2 | 3 | 4 | 4 | 4 | 4 | 4 | 4 | 3 | 4 | 3 |
| Cotton_A_11914_BGI-A2_v1.0 | 38 | 3.2 | Gosto.005G051300-1, Gosto.013G208400-1 | 100.0% | OG0000861 | 3 | 2 | 4 | 4 | 4 | 4 | 4 | 2 | 4 | 2 | 4 | 1 |
| Cotton_A_23939_BGI-A2_v1.0 | 30 | 2.5 | Gosto.001G196400-1 | 95.6% | OG0001667 | 3 | 2 | 1 | 3 | 3 | 3 | 3 | 2 | 3 | 3 | 4 | 1 |
| Cotton_A_28295_BGI-A2_v1.0 | 23 | 1.9 | Gosto.006G027000-1 | 97.9% | OG0002853 | 2 | 2 | 2 | 2 | 2 | 2 | 2 | 2 | 2 | 2 | 4 | 0 |
| Cotton_A_40086_BGI-A2_v1.0 | 22 | 1.8 | Gosto.011G284200-1 | 95.9% | OG0004295 | 2 | 1 | 2 | 2 | 2 | 2 | 2 | 2 | 2 | 2 | 2 | 1 |
| Cotton_A_19720_BGI-A2_v1.0 | 21 | 1.8 | Gosto.013G033400-1 | 100.0% | OG0004345 | 2 | 2 | 2 | 2 | 2 | 2 | 2 | 1 | 1 | 2 | 2 | 1 |
| Cotton_A_00757_BGI-A2_v1.0 | 14 | 1.2 | Gosto.003G217500-1 | 97.7% | OG0006575 | 2 (2) | 1 | 1 | 1 | 1 | 1 | 1 | 1 | 2 (2) | 1 | 1 | 1 |
| Cotton_A_00151_BGI-A2_v1.0 | 12 | 1.0 | Gosto.001G010500-1 | 98.2% | OG0008663 | 1 | 1 | 1 | 1 | 1 | 1 | 1 | 1 | 1 | 1 | 1 | 1 |
| Cotton_A_00108_BGI-A2_v1.0 | 12 | 1.0 | Gosto.001G014500-1 | 97.7% | OG0008676 | 1 | 1 | 1 | 1 | 1 | 1 | 1 | 1 | 1 | 1 | 1 | 1 |
| Cotton_A_19100_BGI-A2_v1.0 | 12 | 1.0 | Gosto.007G179500-1 | 96.0% | OG0012201 | 1 | 1 | 1 | 1 | 1 | 1 | 1 | 1 | 2 (2) | 1 | 1 | 0 |
| Cotton_A_29104_BGI-A2_v1.0 | 12 | 1.0 | Gosto.009G260900-1 | 97.2% | OG0013132 | 1 | 1 | 1 | 1 | 1 | 1 | 1 | 1 | 1 | 1 | 1 | 1 |
| Cotton_A_01472_BGI-A2_v1.0 | 12 | 1.0 | Gosto.012G280100-1 | 95.2% | OG0015696 | 1 | 1 | 1 | 1 | 1 | 1 | 1 | 1 | 1 | 1 | 1 | 1 |
| Cotton_A_25246_BGI-A2_v1.0 | 11 | 0.9 | none | --- | OG0018038 | 1 | 0 | 1 | 1 | 1 | 1 | 1 | 1 | 1 | 1 | 1 | 1 |
| Cotton_A_17591_BGI-A2_v1.0 | 10 | 0.8 | Gosto.004G048300-1 | 98.0% | OG0021151 | 1 | 1 | 1 | 1 | 1 | 0 | 1 | 0 | 1 | 1 | 1 | 1 |

\* candidates derived from Naqvi *et al.* 2017; reference genome is Li *et al.* 2014

### Conclusion

Cotton leaf curl virus is an important cotton pathogen that results in thickening and yellowing of small leaf veins, ultimately leading to the characteristic leaf “curling” phenotype, as well as stunted growth, delayed onset of flowering and/or fruiting, and reductions in yield quantity and quality (Rahman *et al.* 2001; Farooq *et al.* 2015; Rehman *et al.* 2017). Here we report a genome sequence for *Gossypium stocksii,* one of the poorly understood “E-genome” species, which is also a source of CLCuD resistance. This resource provides a new foundation for understanding CLCuD resistance in cotton and represents a new resource for future evolutionary and taxonomic work in this group of cotton species.

## Acknowledgements

We thank the National Science Foundation Plant Genome Research Program (Grant #1339412), Cotton Inc., and the United States Dept. of Agriculture - Agriculture Research Service (Grant #58-6066-6-046) for their financial support. We thank the Iowa State University ResearchIT unit and the BYU Fulton SuperComputer lab for computational resources and support.

